# SETD2 deficiency promotes renal fibrosis through the TGF-β/Smad signaling pathway in the absence of VHL

**DOI:** 10.1101/2022.09.10.507394

**Authors:** Changwei Liu, Xiaoxue Li, Yiwen Zhu, Wenxin Feng, Wei Zhang, Chunxiao Ma, Yue Xu, Liming Gui, Rebiguli Aji, Wei-Qiang Gao, Li Li

## Abstract

Renal fibrosis is the final development pathway and the most common pathological manifestation of chronic kidney disease. An important intrinsic cause of renal fibrosis is epigenetic alterations. SET domain–containing 2 (SETD2) is the sole histone H3K36 trimethyltransferase, catalyzing H3K36 dimethylation to trimethylation. There is evidence that SETD2-mediated epigenetic alterations are implicated in many diseases. However, it is unclear what role SETD2 plays in the development of renal fibrosis. Clinical data indicate that SETD2 is lowly expressed in patients with renal fibrosis. Here, we established genetically engineered mice with SETD2 and VHL deficiency. SETD2 deficiency leads to severe renal fibrosis in VHL-deficient mice. Mechanically, SETD2 maintains the transcriptional level of Smad7, a negative feedback factor of the TGF-β/Smad signaling pathway, thereby preventing the activation of the TGF-β/Smad signaling pathway. Deletion of SETD2 leads to reduced smad7 expression, which results in activation of the TGF-β/Smad signaling pathway and ultimately fibrosis in the absence of VHL. Our findings reveal the role of SETD2-mediated H3K36me3 of Smad7 in regulating the TGF-β/Smad signaling pathway in renal fibrogenesis. Thus, our study provides innovative insights into SETD2 as a potential therapeutic target for the treatment of renal fibrosis.

## Introduction

Chronic kidney disease (CKD) is a progressive disease with no cure and affects approximately 10% of adults worldwide.^1^ Regardless of underlying etiology, renal fibrosis is the final development pathway and the most common pathological manifestation of CKD.^2^ Characterized by deposition of extracellular matrix (ECM), such as α-smooth muscle actin (α-SMA) and fibronectin, renal fibrosis causes tissue scarring and ultimately leads to end-stage of renal disease.^3^ For most people with kidney failure, kidney transplantation or dialysis remains the most common treatment option due to the irreversible nature of the end-stage renal disease. There are, however, several challenges associated with the treatment of end-stage renal disease, including a shortage of kidney donors, a reduction in quality of life, and a high mortality rate after dialysis.^4–7^ So in addition to early prevention, it is crucial to identify and detect new potential targets to prevent renal fibrogenesis.

There is compelling literature suggesting increased expression and activation of transforming growth factor-β (TGF-β/Smad) in human kidney disease.^8,9^ Furthermore, many animal studies have been established to confirm TGF-β/Smad signaling pathway is the primary factor that drives renal fibrosis.^10,11^ TGF-β/Smad signaling pathway acts through a very typical signaling pathway that includes the phosphorylation and activation of Smad2 and Smad3 by the TGF-β/Smad signaling pathway receptor 1 (TGFR1, also known as ALK5), then the phosphorylated Smad2 and Smad3 bind Smad4 to form a complex, which translocates to the nucleus and promotes the transcription of specific genes.^12^ This process is inhibited by the inhibitory Smad (I-Smad), including Smad6 and Smad7. As a negative feedback inhibitor of the TGF-β/Smad signaling pathway, Smad7 recruits E3 ubiquitin ligase SMAD-ubiquitination-regulatory factor 1 (Smurf1) and Smurf2 to degrade TGFRI.^13^ In addition, Smad7 can inhibit the TGF-β/Smad signaling pathway by competing with smad2/3 for bounding to TGFRI to prevent the phosphorylation of Smad2/3.^14^

An important intrinsic cause of CKD is epigenetic alteration.^15^ Epigenetics refers to changes in the transcription and expression of genes rather than changes in the genes themselves, including methylation of DNA, histone modification, and non-coding RNA^16^. There are compelling studies suggesting histone modification plays a key role in CKD and renal fibrogenesis, such as H3K9me2/3, H3K4me1/2/3, and H3K27me3.^17,18^ However, the role of H3K36 methylation in renal fibrosis remains unknown. SET domain– containing 2 (SETD2), the sole histone H3K36 trimethyltransferase, which catalyzes H3K36 dimethylation to trimethylation.^19^ SETD2 has been described to be implicated in a broad range of biological processes, including DNA repair, transcription initiation and elongation, and alternative splicing.^20–22^ It is worth noting that SETD2 loss perturbs the kidney cancer epigenetic landscape to promote metastasis and engenders actionable dependencies on histone chaperone complexes. Our recent studies also reported that SETD2 plays important role in developmental areas and disease occurrence.^23–31^ Notably, SETD2 deficiency accelerates the transition from polycystic kidney disease (PKD) to renal cell carcinoma (RCC) by regulating β-catenin activity.^31^ However, the role of SETD2 in renal fibrosis remains still unknown.

Here, we established a mouse model of renal fibrosis driven by the inactivation of SETD2 and VHL. We found that SETD2 can maintain the transcriptional level of Smad7 through H3K36me3, thus inhibiting the activation of the TGF-β/Smad signaling pathway. SETD2 depletion results in decreased smad7 level and TGF-β/Smad signaling pathway activation which is predictive of fibrosis in the absence of VHL.

## Results

### The mRNA expression of SETD2 is positively correlated with VHL in kidney tissue

To investigate whether SETD2 expression is altered in fibrotic kidneys, we analyzed clinical data from Keenan Research Centre for Biomedical Science and found that reduced expression of SETD2 in the kidney of patients with failure kidneys (Figure 1A).^32^ Meanwhile, considering that VHL deficiency can result in increased vasculature and fibrosis,^33,34^ we explored the relationship between SETD2 and VHL. The mRNA expression of VHL in the kidneys with renal fibrosis is also reduced compared to normal kidneys (Figure 1B). In addition, the mRNA expression of SETD2 is positively correlated with VHL in both failure and normal kidneys (Figure 1C and D).

**Figure 1.**
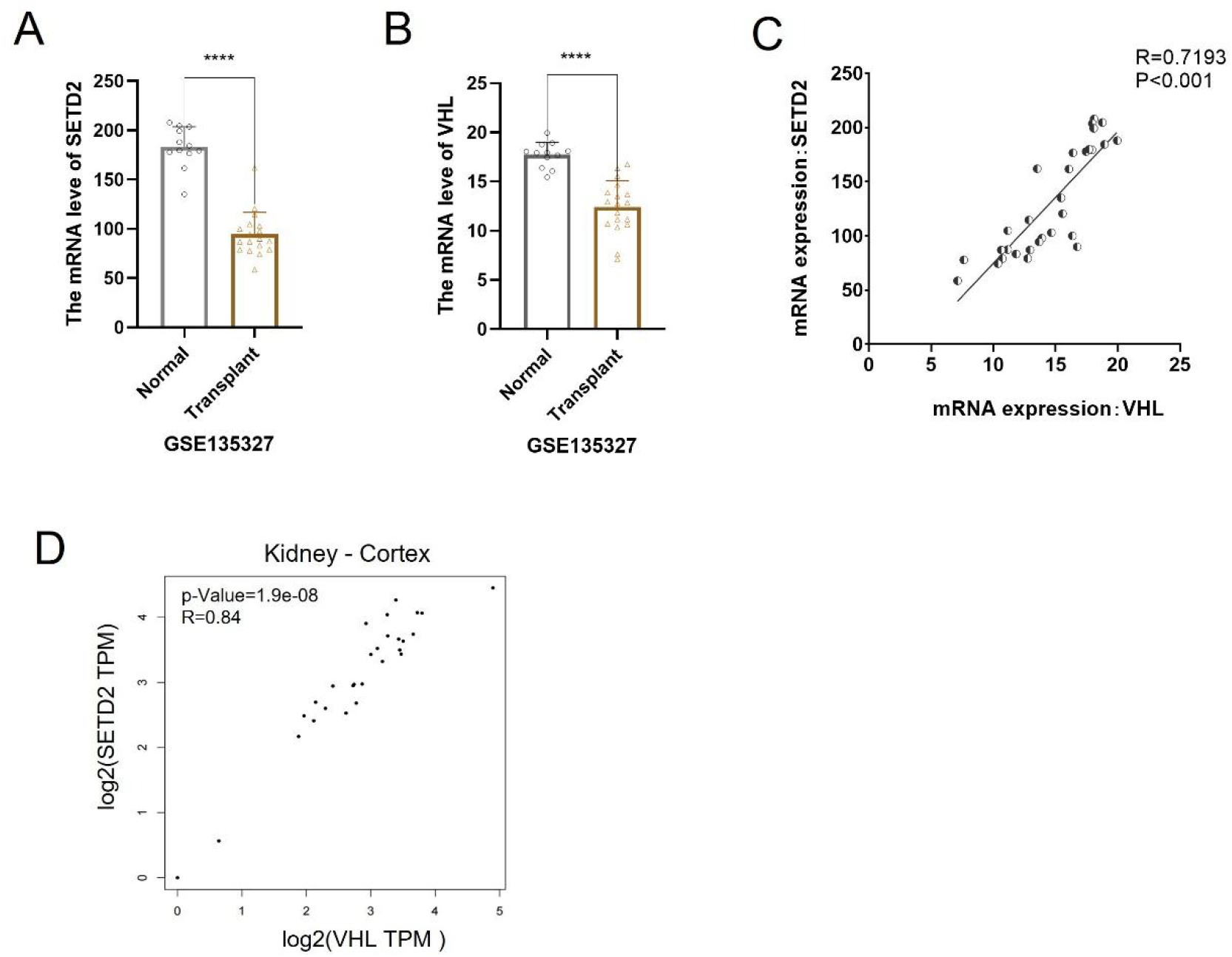
The mRNA expression of SETD2 is positively correlated with VHL in kidney tissue. (A) The mRNA expression level of SETD2 in the kidneys from normal(n=12)and transplant (n=18) (using dataset GSE13527). (B) The mRNA expression level of VHL in the kidneys from normal(n=12) and transplant (n=18) (using dataset GSE13527). (C) The relation of expressions of SETD2 and VHL in normal and transplant kidneys (n=30) (using dataset GSE13527). (D) The relation of expressions of SETD2 and VHL in normal and transplant kidneys from TCGA.

### SETD2 deficiency resulted in severe renal fibrosis in VHL-deficient mice

To explore the potential function of SETD2 in renal fibrosis *in vivo*, we used a transgenic Ksp1.3/Cre mouse line to generate SETD2^fl/fl^ mice and deleted SETD2 in tubular epithelial cells (referred to as SETD2^−KO^ mice).^35^ However, SETD2 inactivation failed to cause renal fibrosis until 30 weeks old. To further investigate whether SETD2 contributes to renal fibrosis in the absence of VHL, we generated Ksp^Cre^; VHL^fl/fl^ mice (hereafter referred to as VHL^-KO^ mice) and Ksp^Cre^; VHL^fl/fl^; SETD2^fl/fl^ mice (hereafter referred as VHL^-KO^; SETD2^−KO^ mice) (Figure 2A). Inactivation of SETD2 significantly reduced the level of H3K36me3 in tubular epithelial cells. The kidneys from VHL^-KO^; Setd2^−KO^ mice appears to be smaller than that of VHL^-KO^ mice and the kidneys of experimental mice appear to be crumpled (Figure 2B). In addition, VHL^-KO^; Setd2^−KO^ mice exhibited a loss of normal renal tubular structure and numerous vesicles. On further histopathological examination, VHL^-KO^; SETD2^-KO^ mice kidneys displayed typical features of renal fibrosis (Figure 2C and D). These results suggested that the absence of SETD2 may lead to the accumulation of collagen and other ECM composition in VHL-deficient kidneys. The levels of COL1A1, aSMA, and FN protein and mRNA in SETD2^-KO^; VHL^-KO^ mice are much higher than that in VHL^-KO^ mice, suggesting that SETD2 deficiency leads to renal fibrosis (Figure 2E and F). Together, SETD2 deficiency results in severe renal fibrosis in VHL-deficient mice.

**Figure 2.**
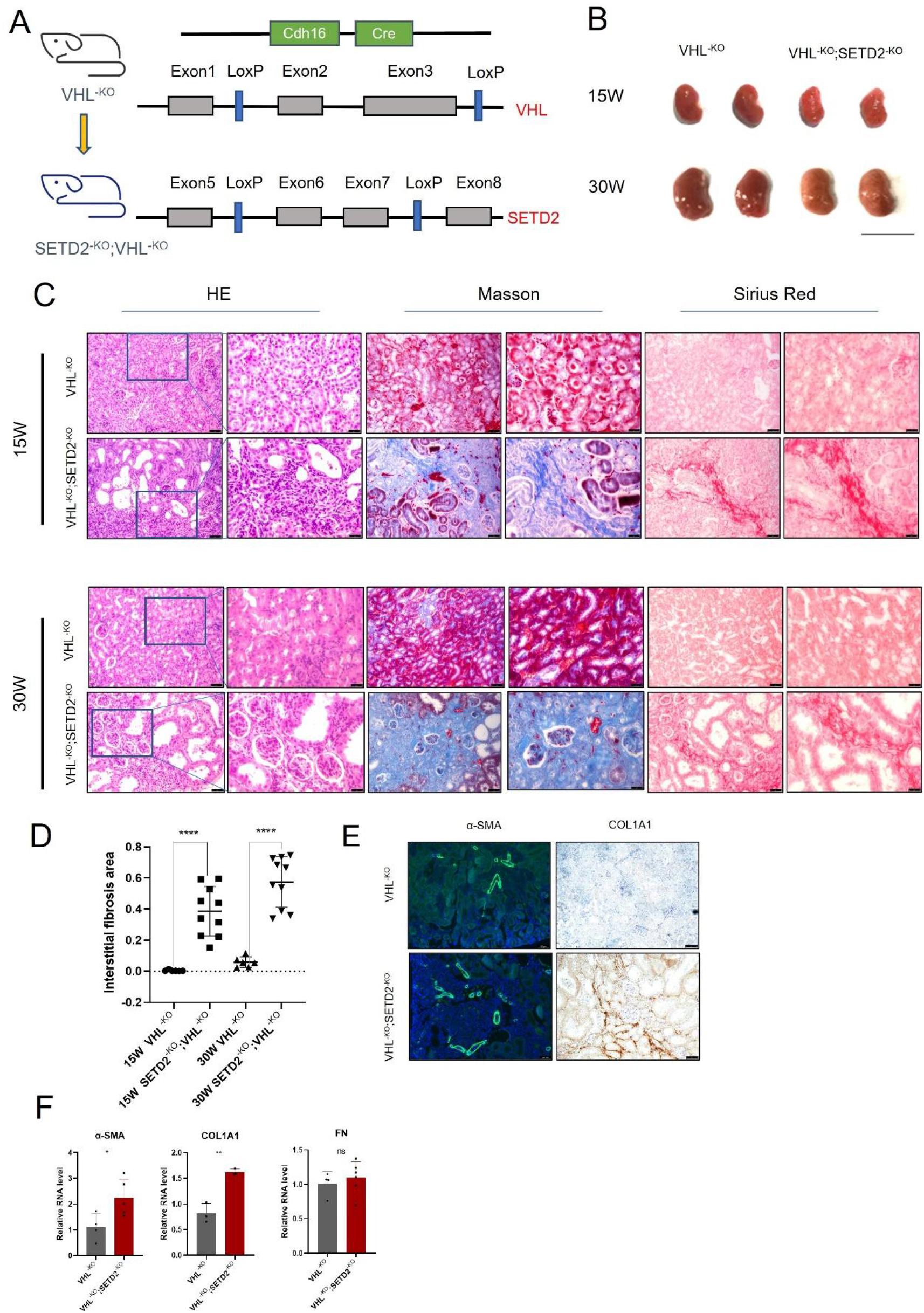
SETD2 deficiency resulted in severe renal fibrosis in VHL-deficient mice. (A) Schematic representation of generating the VHL^-KO^ and VHL^-KO^; SETD2^-KO^ mouse model. (B) Kidney volumes VHL^-KO^ and VHL^-KO^; SETD2^-KO^ mice. Scale bars, 1 cm. (C) and (D)HE, Masson and sirius red staining and statistical results of VHL^-KO^ mice and VHL^-KO^; SETD2^−KO^ mice. Scale bars=50μm. (E) and (F) The protein and mRNA levels of renal fibrosis makers VHL^-KO^ mice and VHL^-KO^; SETD2^−KO^ mice, Scale bars=50μm.

### The inactivation of SETD2 caused hyperactive TGF-β/Smad signaling pathway in VHL-deficient renal tubular epithelial cells

To explore the mechanisms by that SETD2 deficiency promotes renal fibrosis in VHL-deficient renal tubular epithelial cells, we performed RNA-seq using PTECs freshly isolated from 20-week-old VHL^-KO^ and VHL^-KO^; SETD2^-KO^ mice. The overall transcriptome of VHL^-KO^; SETD2^-KO^ PTECs was significantly altered compared to VHL^-KO^ PTECs (Figure 3A). GO analysis and pathway analysis presented that SETD2 inactivation led to a decrease in metabolism-related pathways and significantly enriched lots of genes associated with pathways associated with fibrogenesis in VHL^-KO^; SETD2^-KO^ mice. In addition, GO and pathway analysis showed several pathways were upregulated in VHL^-KO^; SETD2^-KO^ mice compared with VHL^-KO^ mice, including TGF-β signaling pathway, and PI3K signaling pathway (Figure 3B). To better understand the SETD2-mediated signaling pathway, we performed gene set enrichment analysis (GSEA). The data showed that TGF-β/Smad signaling pathway-associated genes were enriched in VHL^-KO^; SETD2^-KO^ mice. To verify the results of the pathway analysis and explore the pivotal pathway in fibrogenesis, qRT-PCR analysis was used to test the expression of important molecules of the above signaling pathways, and we concluded that the TGF-β/Smad signaling pathway was indeed significantly activated in VHL^-KO^; SETD2^-KO^ mice (Figure 3D). TGF-β/Smad signaling pathway plays an important role in renal fibrogenesis and a wide range of studies have established the TGF-β/Smad signaling pathway as the paramount pathogenic factor that drives fibrosis. The activated Smad2 and Smad3 have been verified in VHL^-KO^; SETD2^-KO^ mice by western blotting (WB) (Figure 3E). Furthermore, the data of immunohistochemistry (IHC) also showed that the positive rate of p-Smad3 is much higher in VHL^-KO^; SETD2^-KO^ mice than it’s in VHL^-KO^ mice (Figure 3F). These results indicate that SETD2 can inhibit the TGF-β/Smad signaling pathway transcriptional activity and SETD2 loss leads to p-Smad2 and p-Smad3 activation, which activates TGF-β/Smad signaling and promotes the excessive accumulation of ECM in VHL-deficient kidneys.

**Figure 3.**
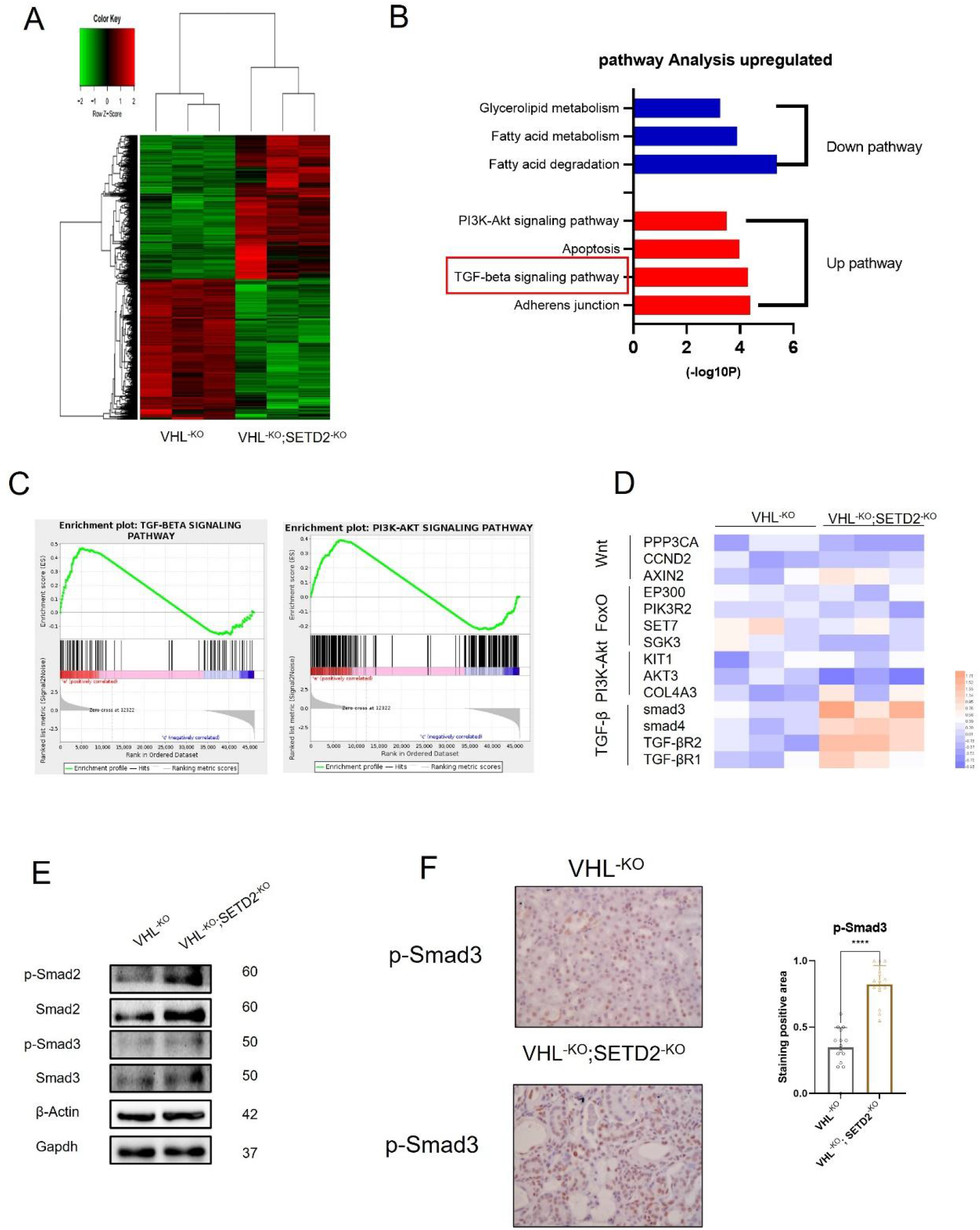
The inactivation of SETD2 caused hyperactive TGF-β/Smad signaling pathway in VHL-deficient renal tubular epithelial cells. (A) Heatmap of genes with significantly different expression in freshly isolated renal tubular epithelial cells based on unsupervised hierarchical agglomerative clustering. (B) Pathway analysis of differentially expressed genes belonging to signaling pathways associated with Setd2 deletion. (C) GSEA enrichment plots of differentially expressed genes associated with Setd2 deletion in VHL deficiency tissue. (D) qRT-PCR analysis of Wnt signaling pathway, TGF-β signaling pathway, FoxO signaling pathway and PI3K signaling pathway. (E) The protein levels of p-Smad2 and p-Smad3 in VHL^-KO^ and VHL^-KO^; SETD2^-KO^ mice. (F) and (G) IHC straining reveal the positive rate of p-Smad3 in VHL-KO and VHL-KO; SETD2-KO mice. Statistical comparisons were made using a two-tailed Student t test. Data are represented as mean ±SEM.

### SETD2 inhibits TGF-β/Smad signaling activation by regulating Smad7 expression

To explore the potential mechanisms of SETD2-dependent epigenetic alterations and to identify genes associated with SETD2 and H3K36me3 on a genome-wide scale, we used chromatin immunoprecipitation sequencing (ChIP-Seq) data to explore possible SETD2-mediated targets. We found that the H3K36me3 antibody occupies Smad7 transcribed region (Figure 4A). To demonstrate the involvement of SETD2 and the H3K36me3 histone marker in Smad7, we verified the occupancy of H3K36me3 in the Smad7 transcribed region using ChIP-qPCR (Figure 4B). These results indicated that SETD2 deletion eliminates H3K36me3 modification in the transcriptional region Smad7. In addition, Smad7 protein expression was reduced (Figure 4C). IHC results showed that SETD2 loss also resulted in the downregulation of Smad7 expression in VHL^-KO^; SETD2^-KO^ PTECS (Figure 4D and E). Smad7 mRNA expression also showed the same results (Figure 4F). To better verify the fact that SETD2 can regulate Smad7 transcription *in vitro*, we utilized renal cell lines and demonstrated that SETD2 depletion leads to a reduction in Smad7 transcript levels, while overexpression of SETD2 leads to a large increase in Smad7 expression (Figure 4G). In addition, clinical data also showed a positive correlation between SETD2 and Smad7 expression (Figure 4H). Together, SETD2 can inhibit TGF-β/Smad signaling activation by regulating Smad7 expression.

**Figure 4.**
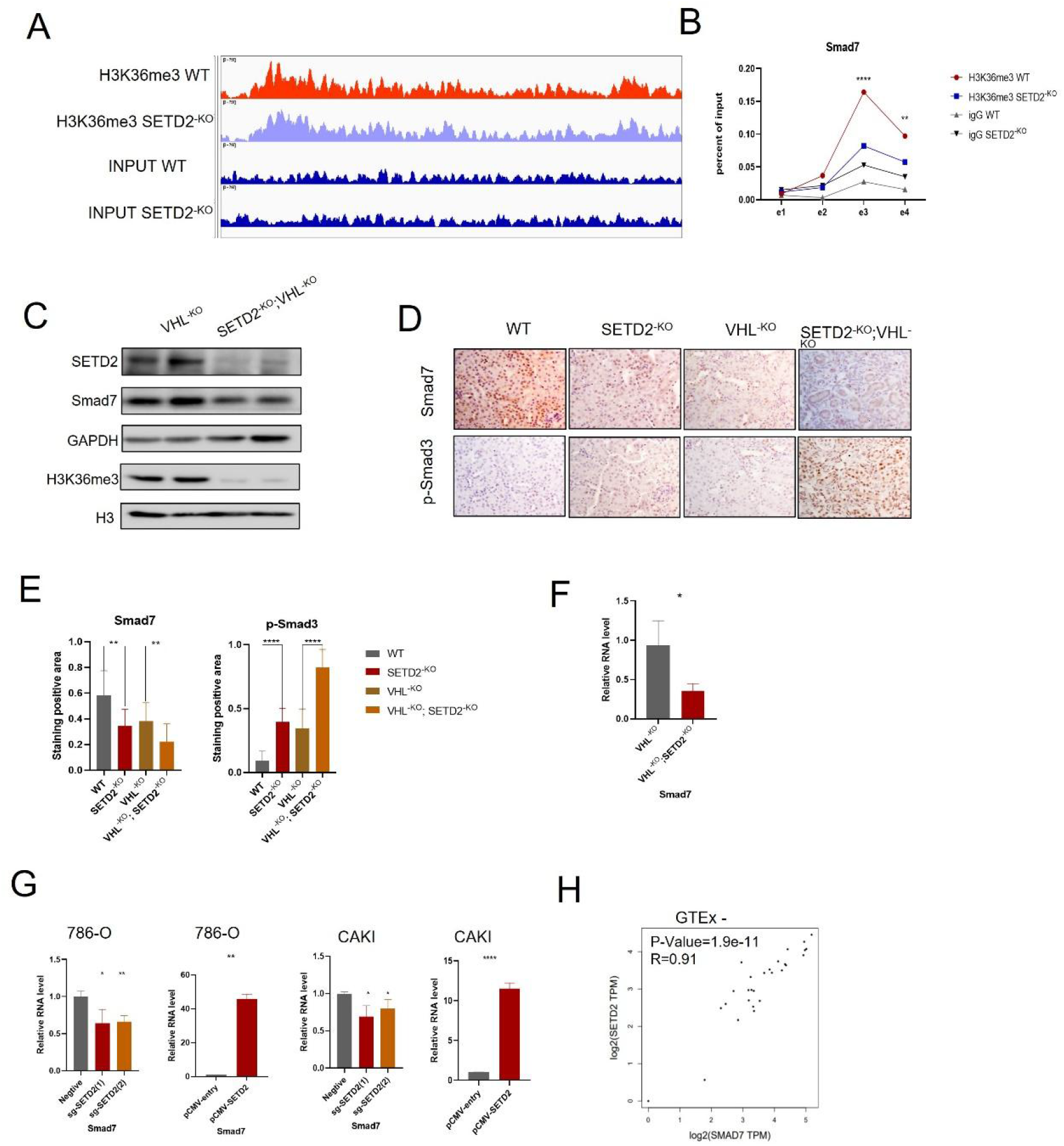
SETD2 inhibits TGF-β/Smad signaling activation by regulating Smad7 expression. (A) Snapshot of H3K36me3 ChIP-Seq signals at the Smad7 gene loci in PTECs. (B) ChIP-qPCR analysis of H3K36me3 occupancy to gene body-retained locus using IgG as the control. (C) Western blot analysis of the Smad7 protein level in VHL^-KO^ and VHL^-KO^; SETD2^-KO^ mice. (D) and (E) IHC straining and statics of Smad7 and p-Smad3 in WT, SETD2^-KO^, VHL^-KO^ and VHL^-KO^; SETD2^-KO^ mice. (F) mRNA levels of Smad7 in VHL^-KO^; SETD2^-KO^ mice. (G) Changes in smad7 mRNA expression in 786-O and CAKI cells after SETD2 knockdown or overexpression. (H) Correlation between SETD2 and Smad7 expression levels in normal kidney tissues. Statistical significance was determined using the Pearson correlation coefficient.

### SETD2 deletion leads to fibrosis through activation of the TGF-β/Smad signaling pathway *in vitro*

To further validate the effect of SETD2 on the TGF-β/Smad signaling pathway and renal fibrosis, we deleted SETD2 through the CRISPR-Cas9 method in HK2. SETD2 loss led to an increase in FN and αSMA levels, demonstrating the deposition of ECM. In addition, SETD2 deletion resulted in increased expression of p-Smad2 and p-Smad3 and decreased expression of Smad7 in HK2 (Figure 5A). Immunofluorescence (IF) results also presented that SETD2 deficiency can result in increased levels of renal fibrosis and the TGF-β/Smad signaling pathway markers (Figure 5B and C). And SIS3, the TGF-β/Smad signaling pathway inhibitor, can relieve the activation of the TGF-β/Smad signaling pathway and fibrosis formation caused by SETD2 deletion (Figure 5D and E). These results indicate that SETD2 deletion leads to fibrogenesis through activation of the TGF-β/Smad signaling pathway *in vitro*.

**Figure 5.**
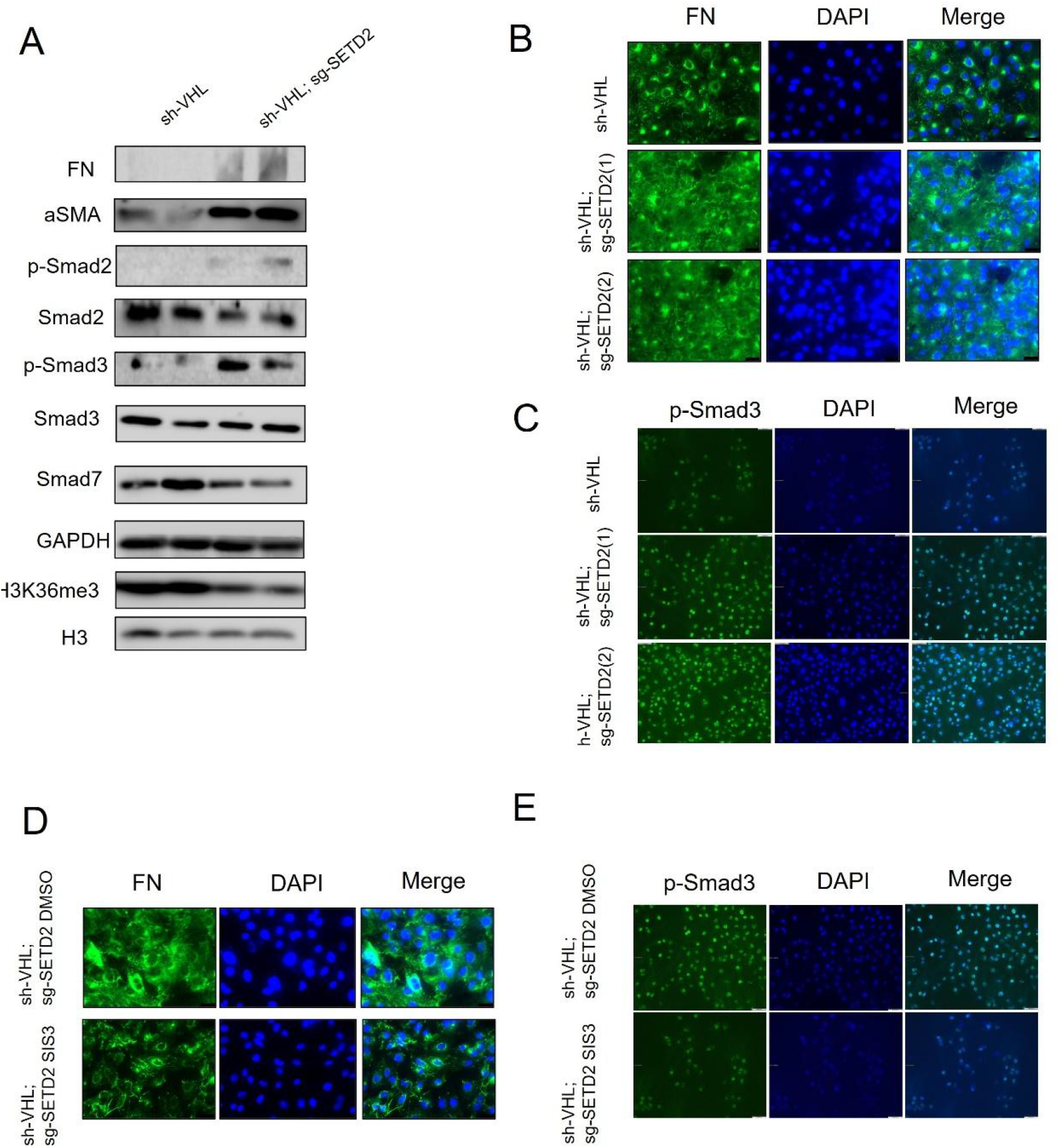
SETD2 deletion leads to fibrosis through activation of the TGF-β/Smad signaling pathway *in vitro*. (A) The expression of indicated proteins in sh-VHL and sh-VHL; sg-SETD2 HK2 cells. (B) and (C) FN and p-Smad3 IF results in sh-VHL and sh-VHL; sg-SETD2 HK2 cells. (C) and (E) FN and p-Smad3 IF results in sh-VHL; sg-SETD2 HK2 cells treated with SIS3 and DMSO.

**Figure 6.**
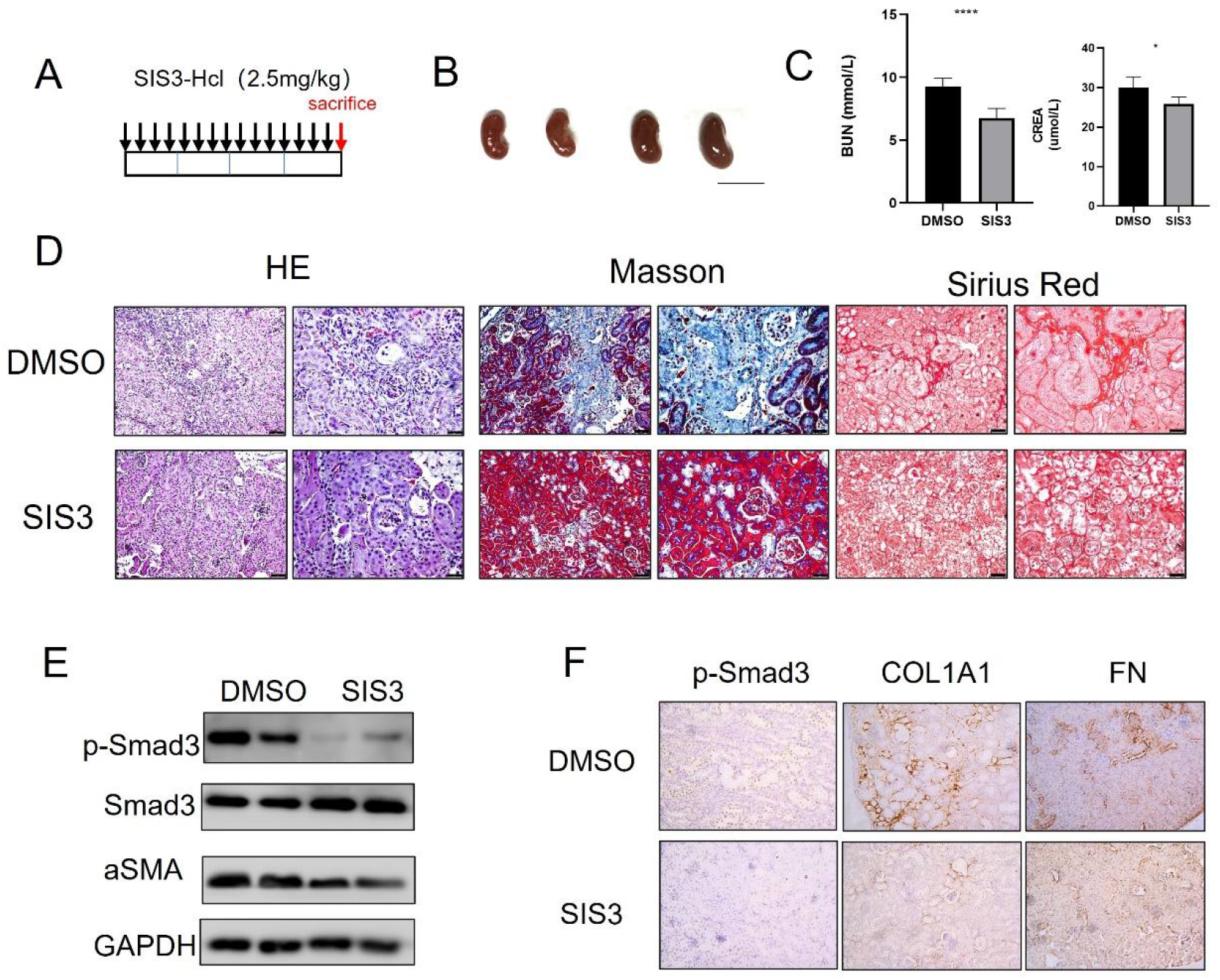
Treatment with TGF-β inhibitor can rescue the renal fibrosis phenotype caused by SETD2 absence. (A) Scheme of treatment were given for each injection. (B) Kidney volumes of VHL^-KO^; SETD2^-KO^ mice treated with SIS3 and DMSO. Scale bars, 1 cm. (C) BUN and crea levels in VHL^-KO^; SETD2^-KO^ mice treated with SIS3 and DMSO. (D) HE, Masson and sirius red staining and statistical results of VHL^-KO^; SETD2^−KO^ mice treated with SIS3 and DMSO. (E) The protein levels of p-Smad3 in VHL^-KO^; SETD2^-KO^ mice treated with SIS3 and DMSO. (F) IHC staining of p-Smad3, Col1a1 and FN in VHL^-KO^; SETD2^-KO^ mice treated with SIS3 and DMSO.

### Treatment with TGF-β inhibitor can rescue the renal fibrosis phenotype caused by SETD2 absence

Considering that SETD2 loss-mediated fibrogenesis is dependent on excessive activation of TGF-β/Smad signaling pathway signaling in HK2, we next investigated whether SIS3 can relieve renal fibrosis in VHL^-KO^; SETD2^-KO^ mice. SIS3 was administered by i.p. injection three times a week and four times a week for four weeks in 12-week-old VHL^-KO^; SETD2^-KO^ mice. We found that the surface of SIS3-treated mice was relatively smooth compared to the control group (Figure 5A and B). BUN and creatinine levels were reduced in the blood of SIS3-treated mice compared to the control group (Figure 5C). In addition, the degree of kidney fibrosis was greatly reduced in SIS3-treated mice compared to the control group (Figure 5D and E). Meanwhile, COL1A1, aSMA, and FN positive staining were reduced in drug-treated mice compared with controls, demonstrating that SIS3 can largely treat the fibrotic phenotype by inhibiting the TGF-β/Smad signaling pathway (Figure 5F and G). Overall, these results suggest that the TGF/β inhibitor rescues the renal fibrosis phenotype caused by SETD2 absence.

## Discussion

In the present study, we demonstrated that SETD2 plays a critical role in the inhibition of the development of kidney fibrosis both *in vitro* and *in vivo*. We have shown that SETD2 deficiency can promote renal fibrosis in VHL-deficient mice by the activation of the TGF-β/Smad signaling pathway. SETD2 maintains the expression level of Smad7 through H3K36me3 and inhibits the TGF-β/Smad signaling pathway. The anti-fibrotic effect of SETD2 provides an innovative insight into SETD2 as a potential therapeutic target for the treatment of renal fibrosis.

SETD2, the sole histone H3K36 trimethyltransferase which catalyzes H3K36 dimethylation to trimethylation, has been reported to participate in diverse biological processes including DNA repair, transcriptional initiation and elongation, and alternative mRNA splicing^20-22^. In addition, our recent studies reported that SETD2 plays important roles in intestinal immunity, suppressing intestinal inflammation, maternal epigenome, genomic imprinting and embryonic development, bone marrow mesenchymal stem cell differentiation, sperm development, and V(D)J recombination in normal lymphocyte development.^23-31^ Moreover, SETD2 loss led to pancreatic carcinogenesis through epigenetic dysregulation of Fbxw7. Based on the polycystic kidney disease (PKD) model caused by the oncogene MYC, SETD2 deficiency accelerates the transition from polycystic kidney disease (PKD) to RCC by regulating β-catenin activity on transcriptional and post-transcriptional levels.

Clinical patient data show that SETD2 is significantly under-expressed in patients with CKD, and what role SETD2 plays in renal fibrosis is still unclear. There is evidence suggesting increased expression and activation of transforming growth factor-β (TGF-β/Smad) in human kidney disease. TGF-β/Smad signaling pathway has important regulatory roles in inflammation, tissue repair, embryonic development and cell growth, differentiation, and immune function. Furthermore, many animal studies have been established to confirm TGF-β/Smad signaling pathway is the primary factor that drives renal fibrosis. Few studies have been conducted between SETD2 and the TGF-β/Smad signaling pathway, especially in the field of renal diseases. Smad7 is a negative feedback inhibitor of the TGF-β/Smad signaling pathway. Smad7 can recruit Smurf1 and Smurf2 to degrade TGFRI. In addition, Smad7 can inhibit the TGF-β/Smad signaling pathway by competing with smad2/3 for bounding to TGFRI to prevent the phosphorylation of Smad2/3. SIS3 is a selective Smad3 inhibitor that attenuates TGF-β1-dependent Smad3 phosphorylation and DNA binding. SIS3 can alleviate many diseases, such as lung cancer^36^. The results that SIS3 can treat fibrosis caused by SETD2 inactivation to some extent prove that the clinical application of SIS3 will be more desirable for these patients with SETD2 reduced expression.

As an important epigenetic regulatory molecule, SETD2 catalyzes not only H3K36 dimethylation to trimethylation but also non-histone substrates, such as α-tubulin, STAT1, and EZH2.^37-39^ In addition, SETD2 also participates in alternative mRNA splicing to affect biological function. In our study, we found that SETD2 affects the expression of Smad7 through H3K36me3 and the TGF-β/Smad signaling pathway. Whether SETD2 can contribute to renal fibrosis through non-histone methylation modifications or interacting with other proteins need to be further investigated.

In summary, we established a mouse model of renal fibrosis driven by deficiency of SETD2 and VHL. SETD2 deficiency resulted in severe renal fibrosis in the absence of VHL, and based on these results, we found that SETD2 deletion can lead to reduced expression of Smad7 by H3K36me3, which activates the TGF-β/Smad signaling pathway and leads to renal fibrosis. In addition, TGF-β inhibitor can rescue the phenotype caused by SETD2 absence. For clinical translation, pharmaceutical investigation of the cross-talks between SETD2 and TGF-β/Smad signaling may provide a potentially promising strategy to prevent renal fibrogenesis in patients.

## Materials and Methods

### Mouse strains

*Setd2*^*fl/fl*^ mice were generated by Shanghai Biomodel Organism Co. The *Ksp*^*Cre*^mice (B6.Cg-Tg (Cdh16-cre) 91Igr/J) and VHL ^*fl/fl*^ were purchased from The Jackson Laboratory. *Setd2*^*fl/fl*^ mice were mated with *Ksp*^*Cre*^ mice to generate *Ksp*^*Cre*^; *Setd2*^*fl/fl*^(*Setd2*^*-KO*^) mice in C57BL/6 background. VHL ^*fl/fl*^ mice were mated with *Ksp*^*Cre*^ mice to generate *Ksp*^*Cre*^; *VHL* ^*fl/fl*^ (*VHL* ^*-KO*^) mice in C57BL/6 background. *Setd2*^*-KO*^ mice were mated with VHL ^*fl/fl*^ mice to generate *Ksp*^*Cre*^; *VHL* ^*fl/fl*^ *Setd2*^*fl/fl*^ (*VHL* ^*-KO*^; *Setd2*^*-KO*^) mice housing under the same condition.

### BUN and creatinine test

Blood was collected from mice using the tail vein blood collection method. Serum was analyzed for BUN and creatinine concentration by Wuhan Servicebio on a Beckman Coulter AU680 analyzer.

### RNA isolation and quantitative RT-qPCR

Total RNA was isolated from cultured cells or fresh samples with Trizol reagent (Invitrogen). cDNA was synthesized by reverse transcription using the Prime Script RT reagent kit (TaKaRa) and subjected to quantitative RT-PCR with Setd2 and target gene primers in the presence of the SYBR Green Realtime PCR Master Mix (Thermo). Relative abundance of mRNA was calculated by normalization to actin-beta or Gapdh mRNA. Data were analyzed from three independent experiments and were shown as the mean ± SEM.

### Western blot analysis and antibodies

Cells were lysed with 100–300 μL RIPA buffer supplemented with protease and phosphatase inhibitors (Millipore). The protein concentration was measured with the BCA Protein Assay (Bio-Rad). From each sample, 20–50 μg of total protein was separated by 8–12% SDS-PAGE gels and transferred onto nitrocellulose membranes (GM). Membranes were blocked in 5% BSA in TBS for 1 hour at room temperature and then incubated with primary antibodies overnight at 4°C, washed in TBS containing 1% Tween20, incubated with an HRP-conjugated secondary antibody for 1 hour at room temperature, and developed by ECL reagent (Thermo). The immunoblots were quantified by Bio-Rad Quantity One version 4.1 software. Primary antibodies against SETD2 (LS-C332416), histone H3 (trimethylK36) (ab9050), Smad7 (ab216428), p-Smad3 (ab52903), COL1A1 (72026S).

### Histology and immunohistochemistry (IHC) Staining

Tissues were fixed in 10% buffered formalin and fixed tissues were sectioned for hematoxylin and eosin (H&E) staining. For IHC staining, paraffin-embedded tissues were deparaffinized, rehydrated, and subjected to a heat-induced epitope retrieval step by treatment with 0.01M sodium citrate (pH 6.0). Endogenous peroxidase activity was blocked with 0.3% (v/v) hydrogen peroxide in distilled water. The sections were then incubated with 0.3% Triton X-100 in PBS (137 mM NaCl, 2.7 mM KCl, 10 mM Na2HPO4, 2 mM KH2PO4, pH 7.4) for 15 minutes, followed by 10% goat serum in PBS for 1 hour. Subsequently, samples were incubated with Primary antibodies, diluted at an appropriate proportion in 1% goat serum for 1 hour at 37°C. After three washes in PBS, sections were incubated with an HRP-conjugated secondary antibody for 1 hour at room temperature. Sections were counterstained with hematoxylin. Three random immunostaining images of each specimen were captured using a Leica DM2500 microscope and analyzed by Image-Pro Plus 6.0 software. Primary antibodies against SETD2 (LS-C332416),

### RNA-seq and analyses

Kidney tissues from mice were harvested around 20-week for RNA preparation. NEB Next Ultra Directional RNA Library Prep Kit for Illumina (New England Biolabs, Ipswich, MA, USA) was used for the construction of sequencing libraries. The libraries were then subjected to Illumina sequencing with paired-ends 2×150 as the sequencing mode. The clean reads were mapped to the mouse genome (assembly GRCm38) using the HISAT2 software. Gene expression levels were estimated using FPKM (fragments per kilobase of exon per million fragments mapped) by StringTie. Gene annotation file was retrieved from Ensembl genome browser 90 databases. To annotate genes with gene ontology (GO) terms and KEGG pathways, ClusterProfiler (R package) was used. The functional enrichment analysis (GO and KEGG) was also performed with ClusterProfiler.

### Statistics

Statistical evaluation was conducted using Student’s t-test. Multiple comparisons were analyzed first by one-way analysis of variance. A significant difference was defined as P <0.05.

### Data availability

The datasets used and/or analysed during the current study are available from the corresponding author on reasonable request. RNA-Seq has been deposited in the Gene Expression Omnibus (GEO) under accession number GEO: deferred public

### Masson and Sirius red

In our study, Masson and Sirius red were performed as described. ^40^

## Acknowledgments

This study was supported by funds from National Natural Science Foundation of China (82073104 to L.L., 81872406 to W.Q.G.), Science and Technology Commission of Shanghai Municipality (19140905500 to L.L., 21JC1404100 to W.Q.G.), KC Wong foundation (to W.Q.G.) and 111 project (no. B21024). The study is also supported by Bio-ID Center, School of Biomedical Engineering, Shanghai Jiao Tong University.

## Author contributions

LL and WQG conceived the concept. LL designed the experiments and revised manuscript. LL and CL prepared the manuscript. CL performed most experiments and analyzed the data. XL and ZW contributed to the animal breeding and identification. WF, WZ, CM, YX, and RA helped with the experiments. All authors edited and approved the final manuscript.

